# Multiple classes of bactericidal antibiotics cause DNA double strand breaks in *Staphylococcus aureus*

**DOI:** 10.1101/2021.03.05.434095

**Authors:** Rebecca S. Clarke, Kam Pou Ha, Andrew M. Edwards

**Affiliations:** MRC Centre for Molecular Bacteriology and Infection, Imperial College London, Armstrong Rd London, SW7 2AZ, UK; Université Paris-Saclay, CEA, CNRS, Institute for Integrative Biology of the Cell (I2BC), 91198, Gif-sur-Yvette, France

**Keywords:** *Staphylococcus*, antibiotic, oxidative stress, DNA, DNA repair

## Abstract

Antibiotics inhibit essential bacterial processes, resulting in arrest of growth and in some cases cell death. Many antibiotics are also reported to trigger endogenous production of reactive oxygen species (ROS), which damage DNA and other macromolecules. However, the type of DNA damage that arises and the mechanisms used by bacteria to repair it are largely unclear. We found that several different classes of antibiotic triggered dose-dependent DNA damage in *Staphylococcus aureus*, including some bacteriostatic drugs. Damage was heterogenous and varied in magnitude between strains. However, antibiotic-triggered DNA damage led to double strand breaks, the processing of which by the RexAB helicase/nuclease complex triggered the SOS response and reduced staphylococcal susceptibility to most of the antibacterials tested. In most cases, DNA DSBs occurred under aerobic but not anaerobic conditions, suggesting a role for ROS. We conclude that DNA double strand breaks are a common occurrence during bacterial exposure to several different antibiotic classes and that repair of this damage by the RexAB complex promotes bacterial survival.

## Introduction

*Staphylococcus aureus* is a common cause of both superficial and invasive infections [1]. Many of these infections, such as infective endocarditis and osteomyelitis can be difficult to treat, requiring lengthy courses of therapy [2,3,4,5,6,7,8,9,10]. Staphylococcal infections are also associated with a high rate of relapse and/or the development of chronic infections, even when the bacteria causing the infection appear to be fully antibiotic susceptible [2,3,4,5,6,7,8,9,10].

There is, therefore, a pressing need to identify new approaches to enhance antibiotic efficacy. To do this, it is important to have a comprehensive understanding of the factors that influence bacterial susceptibility to antibiotics. For example, replication rate has been shown to correlate with susceptibility to several classes of antibiotic [11,12,13]. However, recent evidence suggests that metabolic activity is a better indicator of susceptibility than the replication rate, indicating that metabolism contributes to the bactericidal activity of certain antibacterial drugs [14]. This is because the inhibition of bacterial processes by bactericidal antibiotics leads to metabolic perturbations, which in turn result in the generation of reactive oxygen species (ROS) [15,16,17,18,19,20,21,22]. These highly reactive molecules damage cellular molecules including DNA, lipids and protein and have been proposed to contribute to the lethality of bactericidal antibiotics [15,16,23,24,25,26]. Antibiotic-triggered ROS production has been reported to occur in several different bacteria in response to many different classes of antibiotic [15,16,17,18,19,20,21,22]. However, the magnitude of the damage caused by antibiotic-triggered ROS production and the degree to which these radicals contribute to bacterial killing is unclear [27,28,29]. This issue is worth resolving because a greater understanding of the mechanisms by which bacteria repair the damage caused by ROS may help to identify new therapeutics that enhance antibiotic activity. For example, we have shown previously that the combination antibiotic co-trimoxazole (trimethoprim plus sulphmethoxazole) caused DNA double strand breaks (DSB) and induction of the SOS DNA repair response in an oxygen-dependent manner in the major human pathogen Staphylococcus aureus [30]. This DNA damage was lethal if not repaired, resulting in 50-5000-fold greater reduction in CFU counts of a mutant defective for DSB repair (*rexB*::Tn) relative to wild type *S. aureus*. As such, inhibitors of RexAB would be expected to significantly enhance the bactericidal activity of co-trimoxazole against *S. aureus*.

To understand whether our findings with co-trimoxazole were applicable to other antibacterial drugs, we undertook a comprehensive analysis of multiple classes of antibiotics. This revealed that most antibiotics cause DNA damage in *S. aureus* under aerobic conditions, which results in DNA DSBs, since mutants lacking DNA DSB repair complex RexAB were more susceptible to antibiotic killing. Novel inhibitors that block staphylococcal DNA repair complex RexAB would, therefore, be expected to sensitise *S. aureus* to the bactericidal activity of multiple antibiotics.

## Results

### Multiple classes of antibiotics cause DNA damage in *S. aureus*

DNA damage in most bacteria, including *S. aureus*, triggers activation of the SOS response, which leads to the transcription of genes whose products contribute to DNA repair [30,31,32,33,34]. These genes include *recA*, which encodes the RecA protein required for homologous recombination and, together with LexA, is a key regulator of the SOS response [31,32,33].

To determine whether antibiotics caused DNA damage in *S. aureus*, we used a *PrecA-gfp* reporter system in two distinct genetic backgrounds: SH1000, a methicillin-sensitive *S. aureus* (MSSA) strain, and JE2, a community-associated methicillin-resistant *S. aureus* (CA-MRSA) strain of the USA300 lineage [30,34]. Co-trimoxazole was included as a control, since we have shown previously it triggers the SOS response in *S. aureus* [23].

These strains were then exposed to various classes of clinically relevant antibiotics across a range of concentrations that partially inhibited growth (Fig. S1). These included both bactericidal (co-trimoxazole, ciprofloxacin, nitrofurantoin, oxacillin, daptomycin, gentamicin) and bacteriostatic (chloramphenicol, linezolid) drugs.

As expected, we found that co-trimoxazole, ciprofloxacin, nitrofurantoin and oxacillin triggered SOS induction in both the SH1000 and JE2 strains (Fig. 1A,B,C,D,) [30,34,35,36]. DNA damage was also apparent during exposure to the bactericidal lipopeptide antibiotic daptomycin and the bacteriostatic drugs chloramphenicol and linezolid, albeit to varying degrees and with some differences between the two strains (Fig. 1E,F,G). However, we did not detect any induction of the SOS response during bacterial exposure to gentamicin at any of the concentrations used (Fig. 1H). Taken together, these data indicated that most clinically relevant classes of antibiotics, including bacteriostatic agents caused DNA damage in *S. aureus*.

**Figure 1.**
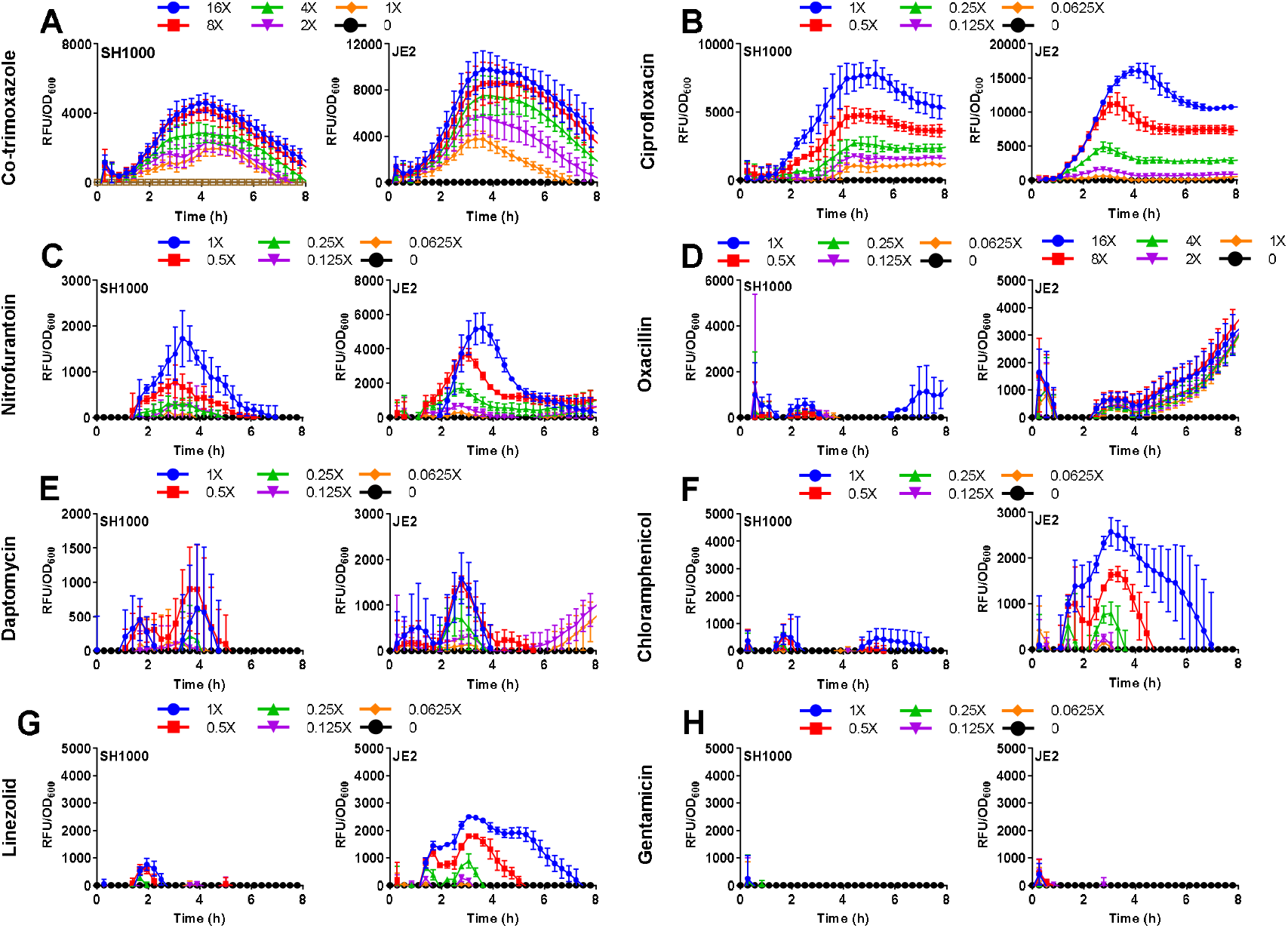
Induction of the SOS response in *S. aureus* SH1000 and JE2 by diverse classes of antibiotics. **(A-H)** Induction of SOS measured by GFP expression driven from a *PrecA-gfp* reporter construct upon exposure to a range of concentrations of various antibiotics. Concentrations were chosen based on their ability to cause growth inhibition and represent multiples of the MIC of the individual strain as indicated in the key above each graph. GFP fluorescence was normalised to OD_600_ to determine induction of SOS relative to cell density. Data represent the mean of 3 independent experiments (n = 3). Representative OD_600_ measurements alone are shown in Fig. S1. Error bars represent standard deviation of the mean.

### SOS induction is partly due to processing of DNA double strand breaks by the RexAB helicase/nuclease complex

We have shown previously that induction of the SOS response by co-trimoxazole is largely due to the processing of DNA double strand breaks (DSBs) by the AddAB-family RexAB nuclease/helicase complex [30,34,37]. Therefore, we determined whether SOS induction by other classes of antibiotics was also due to RexAB-mediated processing of DNA DSBs. As before, co-trimoxazole was included in these assays as a control.

To do this, we compared GFP fluorescence from wild type *S. aureus* JE2 and a *rexB*::Tn mutant defective for RexAB, both of which contained the *PrecA-gfp* reporter system, during exposure to the same panel of antibiotics as described for Figure 1 [30,34,38,39]. As expected from our previous work, we found that the lack of RexAB reduced recA induction relative to the wild type during exposure to co-trimoxazole (Fig. 2A) [30]. We also observed reduced recA expression in the *rexB*::Tn mutant relative to the wild type during exposure to the quinolone antibiotic ciprofloxacin, which is known to cause DNA DSBs (Fig. 2B) [40].

**Figure 2.**
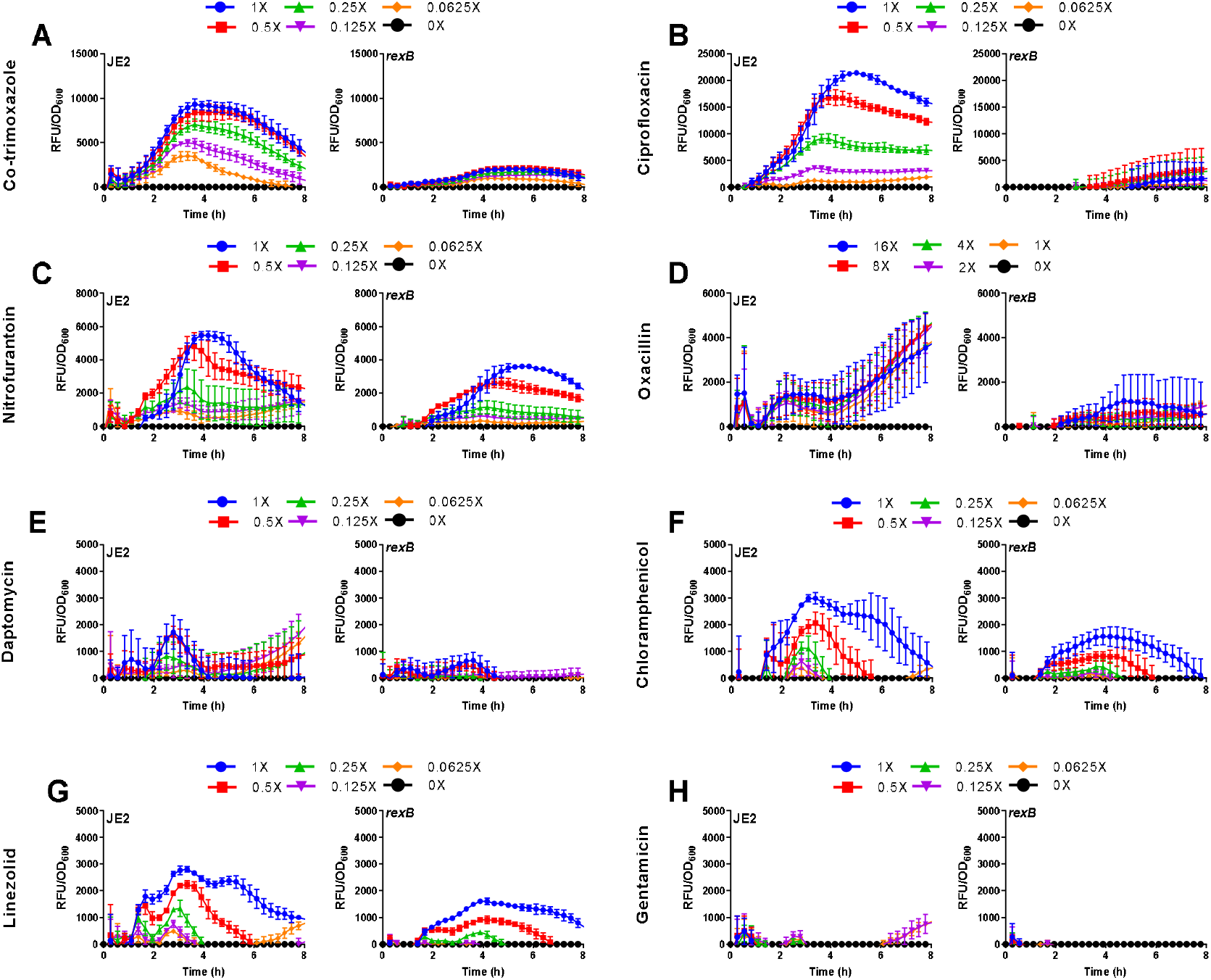
RexAB is required for maximal induction of the SOS response during exposure to antibiotics. **(A-H)** Induction of SOS response of JE2 wild-type and *rexB* mutant measured by GFP expression upon exposure to a range of sub-lethal concentrations of antibiotics. Concentrations of antibiotic are labelled by multiple of the MIC of the wild-type strain. GFP fluorescence was normalised by OD_600_ to determine induction of SOS relative to cell density (n = 3). Representative OD_600_ measurements alone are shown in Fig. S1. Error bars represent standard deviation of the mean.

For nitrofurantoin, oxacillin, daptomycin, chloramphenicol and linezolid, we also observed lower levels of SOS induction in the *rexB*::Tn mutant relative to the wild type, although the differences between strains was not as profound as for co-trimoxazole and ciprofloxacin (Fig. 2C-H). As expected from previous data (Fig. 1G), no *recA* induction was observed from either wild type or *rexB*::Tn mutant during exposure to gentamicin (Fig. 2G). Therefore, as for co-trimoxazole, RexAB is required for maximal induction of SOS in response to DNA damage caused by several clinically relevant antibiotics, indicating that these drugs cause DNA DSBs in *S. aureus*.

### DNA DSB repair reduces bacterial susceptibility to several classes of antibiotics

The requirement of RexAB for maximal induction of the SOS response indicated that exposure to most antibiotics caused DNA DSBs [30,34]. Since DSBs are lethal if not repaired, we hypothesised that mutants defective for RexAB would be more susceptible than wild type strains to those antibiotics that triggered the SOS response.

To test this, we determined the minimum inhibitory concentration (MIC) of each antibiotic for wild type *S. aureus* SH1000 and JE2, and associated *rexB*::Tn mutants (Table 1). The *rexB* mutants in both JE2 and SH1000 strains were ≥2-fold more susceptible to 7 of the 8 antibiotics tested conditions (Table 1). Importantly, the absence of RexAB increased the susceptibility of the MRSA strain JE2 to both oxacillin and ciprofloxacin 4-fold, despite this strain being resistant to both antibiotics.

**Table 1.**
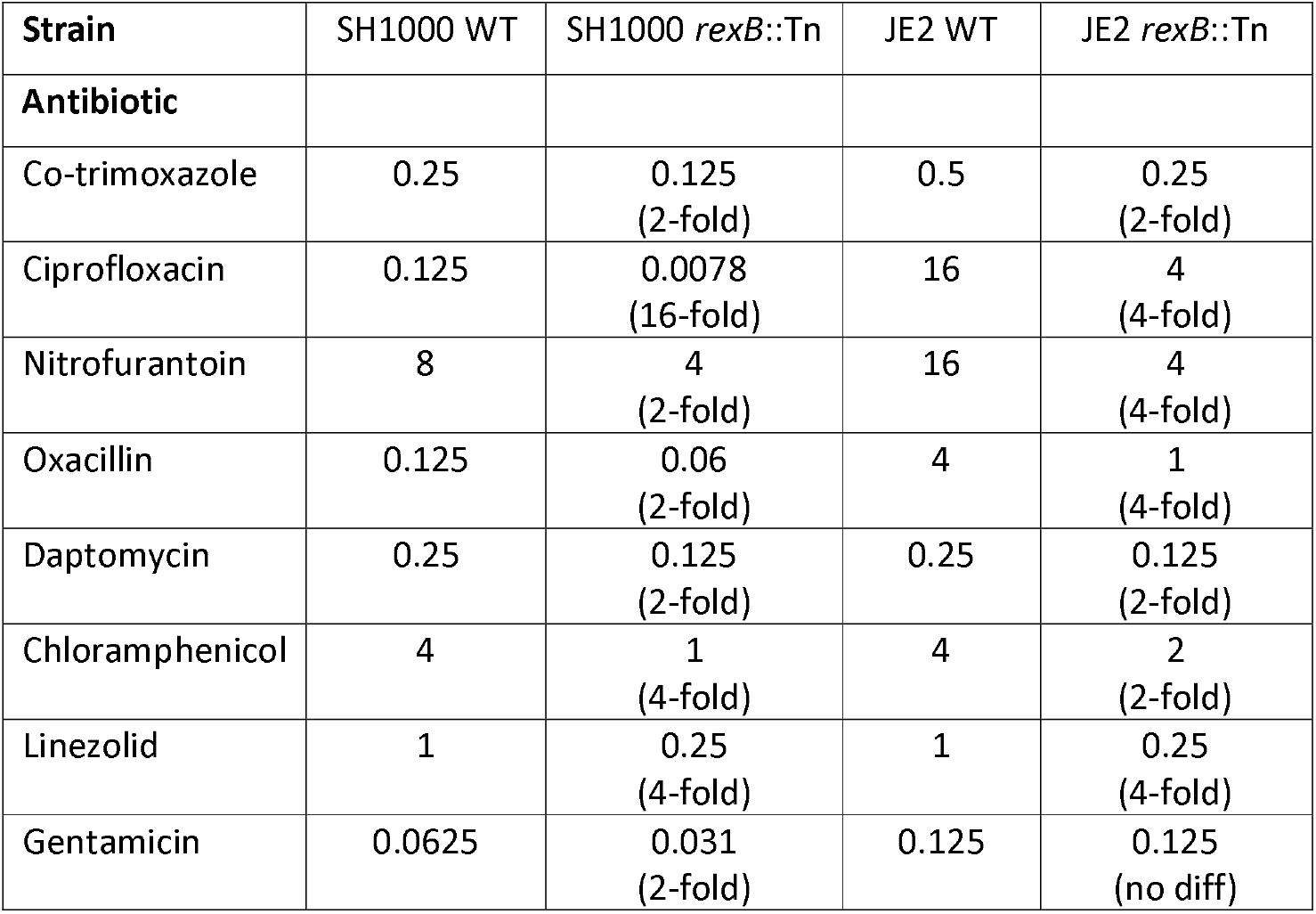
MIC values (μg ml^−1^) of *S. aureus* WT and *rexB* mutant in SH1000 and JE2 backgrounds for various antibiotics (n ≥ 3; median MIC is shown). The fold reduction in MIC of the *rexB*::Tn mutants relative to the wild type are also shown.

| Strain | SH1000 WT | SH1000 *rexB::Tn* | JE2 WT | JE2 *rexB::Tn* |
| --- | --- | --- | --- | --- |
| Antibiotic |  |  |  |  |
| Co-trimoxazole | 0.25 | 0.125<br>(2-fold) | 0.5 | 0.25<br>(2-fold) |
| Ciprofloxacin | 0.125 | 0.0078<br>(16-fold) | 16 | 4<br>(4-fold) |
| Nitrofurantoin | 8 | 4<br>(2-fold) | 16 | 4<br>(4-fold) |
| Oxacillin | 0.125 | 0.06<br>(2-fold) | 4 | 1<br>(4-fold) |
| Daptomycin | 0.25 | 0.125<br>(2-fold) | 0.25 | 0.125<br>(2-fold) |
| Chloramphenicol | 4 | 1<br>(4-fold) | 4 | 2<br>(2-fold) |
| Linezolid | 1 | 0.25<br>(4-fold) | 1 | 0.25<br>(4-fold) |
| Gentamicin | 0.0625 | 0.031<br>(2-fold) | 0.125 | 0.125<br>(no diff) |

The one exception was gentamicin, where the SH1000 *rexB*::Tn mutant was 2-fold more susceptible to the antibiotic, but the JE2 *rexB*::Tn mutant had the same MIC as the wild type strain, in keeping with the fact that this antibiotic did not trigger the SOS response under the conditions tested (Table 1). Taken together, the MIC data provide additional evidence that most antibiotics cause DNA DSBs in *S. aureus*.

### DNA DSB repair promotes staphylococcal tolerance of several classes of antibiotics

We have shown previously that DNA DSB repair by RexAB enables staphylococcal tolerance of the combination antibiotic co-trimoxazole [30]. Since most of the other antibiotics we examined also caused DNA DSBs, leading to increased susceptibility of *rexB*::Tn mutants in MIC measurements, we hypothesised that RexAB would also contribute to bacterial survival during exposure to a supra-MIC concentration of these other anti-bacterial drugs.

To test this we exposed wild type *S. aureus* SH1000 and JE2, and associated *rexB*::Tn mutants to 10X the MIC of the wild type of each of the antibiotics used in previous assays and measured survival after 8 h incubation at 37 C in an aerobic atmosphere (Fig. 3). Similar to the MIC assays, 6 of 8 antibiotics tested were more active against the *rexB*::Tn mutant relative to wild type bacteria, resulting in lower survival of the DNA repair defective strains (Fig. 3). The two antibiotics where there was no difference in survival between wild type and *rexB*::Tn mutants were linezolid and chloramphenicol, which are both bacteriostatic and did not reduce CFU counts of any of the strains (Fig. 3F,G). The remaining 6 antibiotics (co-trimoxazole, ciprofloxacin, oxacillin, nitrofurantoin, daptomycin, gentamicin), all of which are classified as bactericidal, caused significantly greater decreases in CFU counts of the *rexB*::Tn mutants relative to wild type bacteria (Fig. 3A,B,C,D,E,H). The increased susceptibility of the *rexB*::Tn mutants resulted in 5-500-fold greater reductions in CFU counts compared to wild type cells after 8h exposure to the 6 bactericidal antibiotics (Fig. 3A,B,C,D,E,H).

**Figure 3.**
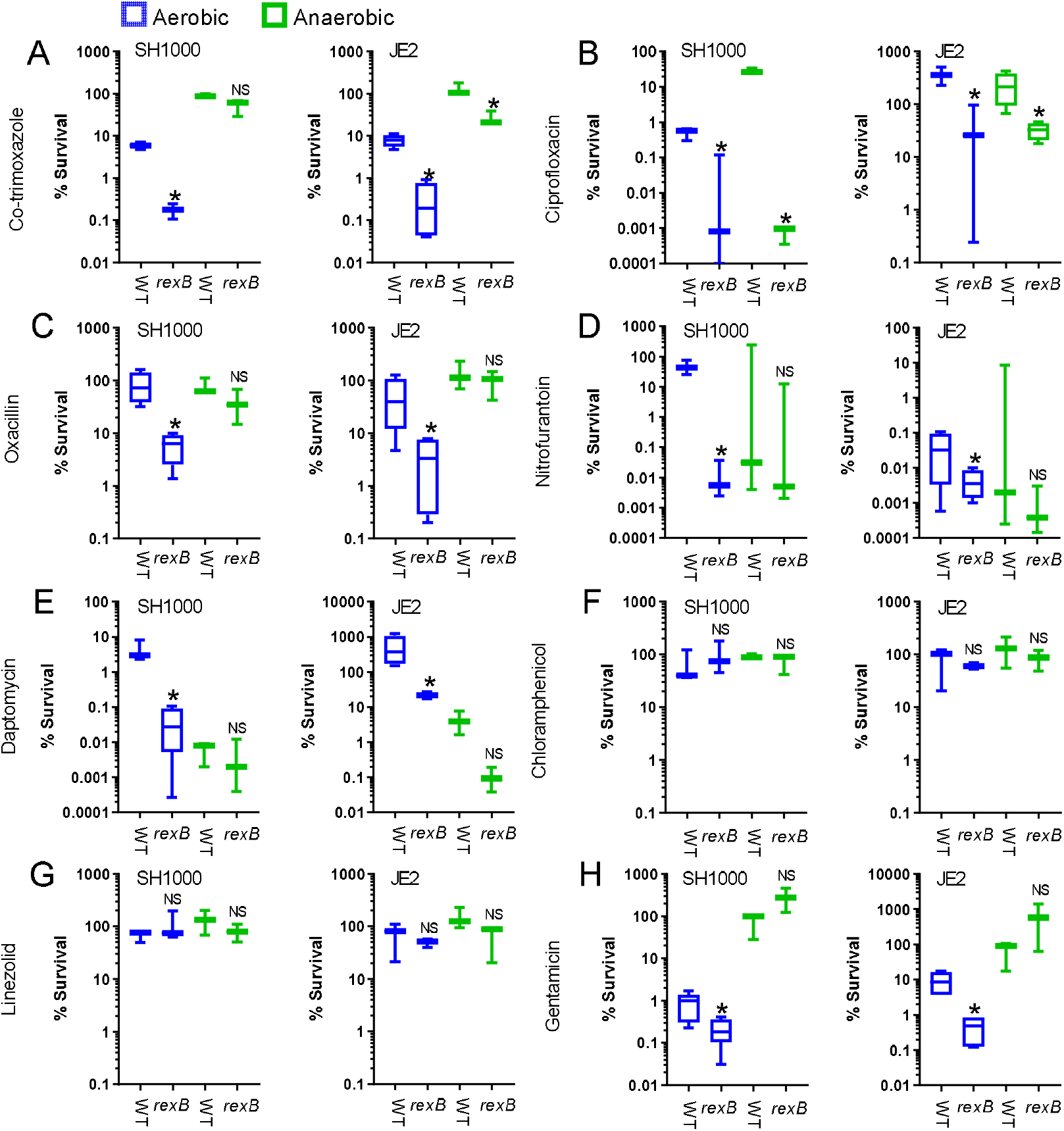
Lack of effective DNA repair increases the killing of *S. aureus* by bactericidal antibiotics under aerobic conditions. **(A-H)** Survival of *S. aureus* WT and *rexB* mutant in SH1000 and JE2 backgrounds after 8 h incubation at 37°C in TSB supplemented with 10x MIC. Survival was assessed under aerobic (blue) or anaerobic (green) conditions (n = 3). Data were analysed by one-way ANOVA (* = P < 0.05) and presented as a box and whisker plot and presented as a box and whisker plot with error bars showing the full data range.

To test whether DNA DSBs caused by bactericidal antibiotics were due to endogenous ROS production, we repeated bactericidal activity assays under anaerobic conditions. As reported previously, co-trimoxazole lost bactericidal activity in the absence of oxygen, as did ciprofloxacin and gentamicin, the latter due to the reduction in membrane potential in the absence of oxygen [30,42] (Fig. 3A,B,H). The bactericidal antibiotics nitrofurantoin and daptomcyin retained bactericidal activity under anaerobic conditions, but oxacillin did not (Fig. 3C,D,E). However, by contrast to aerobic conditions, nitrofurantoin and daptomycin were equally bactericidal against wild type and *rexB*::Tn mutants, suggesting that they caused DNA damage in a ROS dependent manner (Fig. 3).

The finding that loss of RexAB resulted in increased killing of both the SH1000 MSSA and JE2 MRSA strain by the frontline anti-staphylococcal penicillin oxacillin was particularly noteworthy because this indicated a mechanism by which MRSA strains could be re-sensitised to the antibiotic. Therefore, we repeated this assay and included mutants complemented with the *rexB*A operon [34] (Fig. 4). As expected, the *rexB*::Tn mutants were more susceptible to killing by oxacillin than wild type bacteria under aerobic but not anaerobic conditions. Complementation of mutations with plasmids containing the *rexB*A operon, but not the plasmid alone, restored survival to wild type levels, confirming the role of RexAB in staphylococcal tolerance of oxacillin (Fig. 4).

**Figure 4.**
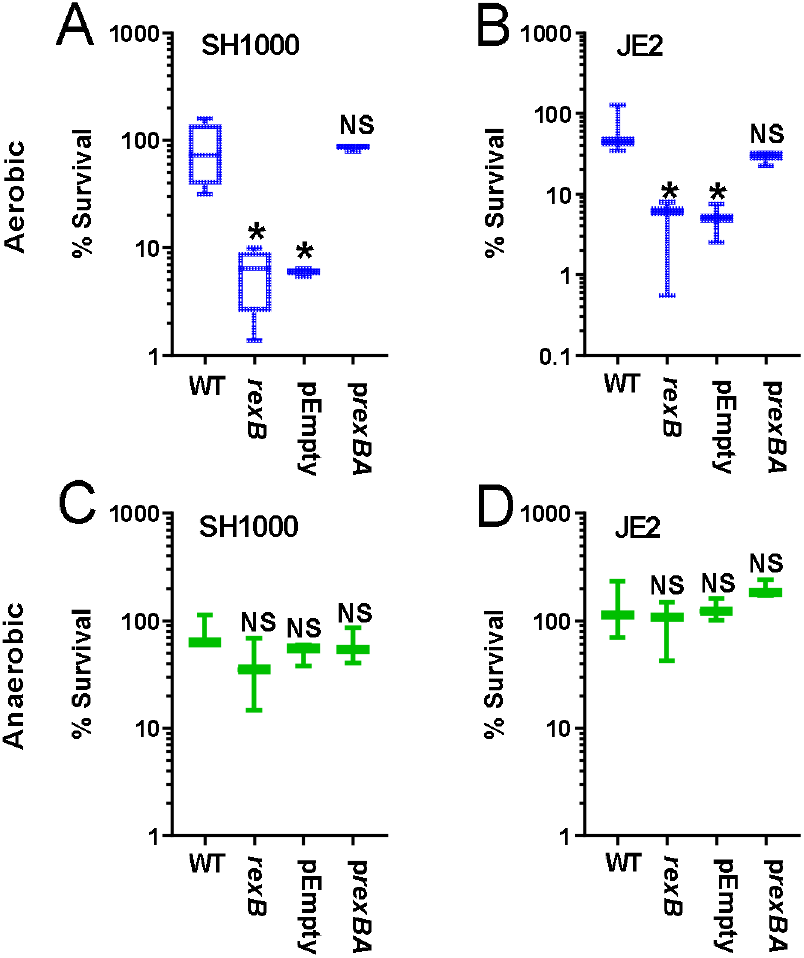
Complementation of the *rexB*::Tn mutant restores tolerance to oxacillin in *S. aureus* SH1000 and JE2. **(A-D)** Survival of *S. aureus* WT and *rexB* mutant in SH1000 and JE2 backgrounds after 8 h incubation at 37°C in TSB supplemented with 10x MIC. Survival was assessed under aerobic (blue) or anaerobic (green) conditions (n = 3). Data were analysed by one-way ANOVA (* = P < 0.05) and presented as a box and whisker plot with error bars showing the full data range.

Taken together, these findings support previous work by showing that several diverse classes of antibiotics cause DNA damage under aerobic conditions, most likely via ROS generation [30]. The data presented here extend these previous observations by demonstrating that antibiotic-mediated DNA damage leads to DNA DSBs and that repair of these DSBs by the RexAB helicase/nuclease complex promotes bacterial survival during exposure to several different classes of antibiotic.

## Discussion

Staphylococcal infections are associated with high rates of treatment failure, even in the case of apparently drug-susceptible strains [1,2,3,4,5,6,7,8,9,10]. This, together with the threat posed by multi drug resistant MRSA strains, necessitates a greater understanding of how antibiotics function and the identification of opportunities to improve their efficacy [9].

There is compelling evidence that diverse antibiotics trigger metabolic perturbations in bacteria that lead to endogenous ROS production under aerobic conditions [14,15,16,17,18,19,20,21,22,23]. However, the consequences of this for bacterial viability remain a matter of debate [27,28,29]. This is important because if endogenous ROS production is a common property of antibiotics then it could be exploited to enhance treatment outcomes, for example by designing inhibitors of bacterial processes that detoxify ROS or repair the damage it causes [30,34,43].

To understand whether antibiotics cause ROS-mediated damage in *S. aureus*, we focussed on the degree to which antibiotic exposure resulted in bacterial DNA damage, since nucleic acids are frequently attacked by endogenous ROS, and the consequences of that damage for bacterial survival [24,25,26,30,34].

Using a *PrecA-gfp* reporter assay we observed that the SOS response in *S. aureus* was triggered by several different classes of antibiotics, indicative of DNA damage. Whilst this was expected for DNA-targeting antibiotics such as the fluoroquinolone ciprofloxacin [40], SOS induction also occurred with antibiotics that do not directly target bacterial DNA such as oxacillin, daptomycin and linezolid. It has been shown that certain β-lactam antibiotics induce the SOS response in *E. coli via* the DpiBA two-component system rather than via DNA damage [44]. Although this mechanism has been hypothesised for *S. aureus*, our data show that *S. aureus rexB*A mutants were more susceptible to killing by oxacillin, demonstrating that DNA damage does occur during exposure to this β-lactam antibiotic and that this is at least partially responsible for triggering the SOS response. Our findings are in keeping with work showing that β-lactam antibiotics trigger endogenous ROS production via elevated TCA cycle activity in response to cell wall damage, leading to increased mutation rate [45].

Whilst still a controversial topic, there is increasing evidence that many classes of antibiotics trigger the endogenous production of ROS. However, the degree to which these ROS contribute to bactericidal a ctivity is less clear. Our data provide evidence that many antibiotics cause DNA damage, presumably via ROS since *rexB*::Tn mutants were as susceptible to most antibiotics under anaerobic conditions. However, this DNA damage appears to be largely tolerated by wild type bacteria via RexAB-mediated processing of DSBs, which triggers the SOS response to facilitate repair via homologous recombination.

The production of ROS by bactericidal but not bacteriostatic antibiotics has been proposed to explain their differences in lethality. However, we observed SOS induction during exposure of *S. aureus* to the bacteriostatic antibiotics linezolid and chloramphenicol, but not the bactericidal antibiotic gentamicin. Surprisingly however, despite DNA damage, the absence of RexAB did not sensitise *S. aureus rexB*::Tn mutants to linezolid or chloramphenicol but did increase susceptibility to gentamicin. This may be explained by differences in the type of DNA damage caused by each of the antibiotics. Whilst several different types of DNA damage trigger SOS, only those leading to DSBs would be expected to promote susceptibility of the *rexB*A mutant [37,45,46,47]. As such, it is possible that bactericidal antibiotics trigger the potentially lethal DNA DSBs, whilst bacteriostatic antibiotics trigger non-lethal types of DNA damage. In keeping with this hypothesis, the absence of RexAB had only a small effect on SOS induction in *S. aureus* caused by linezolid or chloramphenicol. It is unclear why gentamicin did not trigger SOS during antibiotic exposure since it appeared to cause DNA DSBs in the antibiotic tolerance assays, but it may be the case that high concentrations of the antibiotic are needed for DNA damage. Combined, our data indicates differences between antibiotics in the degree of DNA damage caused, as well as the time required to cause damage and these differences may explain some of the debate around the contribution of ROS to antibiotic-mediated killing. However, the data clearly demonstrate that DNA DSBs are a common consequence of the exposure of *S. aureus* to several different classes of antibiotics and that an inability to repair those DSBs increase bacterial susceptibility to the antibacterial drugs. These findings are similar to those reported for E. coli, where mutants defective for DNA DSB repair (defective for *recB* or *recC*) were more susceptible than the wild type to at least 8 different antibiotics [48]. Crucially, we found that disruption of DNA DSB repair restored quinolone susceptibility in an otherwise resistant strain of *S. aureus*, which is also similar to what has been seen in *E. coli* and *Klebsiella pneumoniae* [49]. We also found that an inability to repair DSBs restored oxacillin susceptibility in the JE2 MRSA strain, although it remains to be seen if this finding is applicable to other MRSA strains.

The identification of RexAB as important for staphylococcal survival during exposure to several different antibiotics, including the re-sensitisation of resistant strains to some antibiotics, makes this complex a potential target for novel therapeutics, particularly as the lack of RexAB homologues in eukaryotes reduces the likelihood of host toxicity [37,43,50,51]. Inhibitors of RexAB would be expected to enhance the bactericidal activity of several different classes of antibiotic, as well as reduce the induction of the mutagenic SOS response, which is associated with the emergence of antibiotic resistance and mutants that can resist host immune defences [52,53]. We have also recently shown that DNA DSB repair is important for staphylococcal resistance to host immune defences, in keeping with similar findings with several other bacterial pathogens, providing an additional potential benefit of targeting this complex [34,54,55,56,57].

In summary, staphylococcal DNA is damaged by several classes of bactericidal antibiotics resulting in DSBs that are processed by RexAB and trigger the SOS response for repair. Therefore, RexAB is a potential target for novel therapeutics that sensitise *S. aureus* to antibiotics.

## Materials and Methods

### Bacterial strains and culture conditions

The bacterial strains used in this study are listed in

Table 1. *S. aureus* was cultured in Tryptic Soy Broth (TSB) or Muller Hinton Broth (MHB) to stationary phase (18 h) at 37 °C, with shaking (180 rpm). Media were supplemented with antibiotics as required.

**Table 1.**
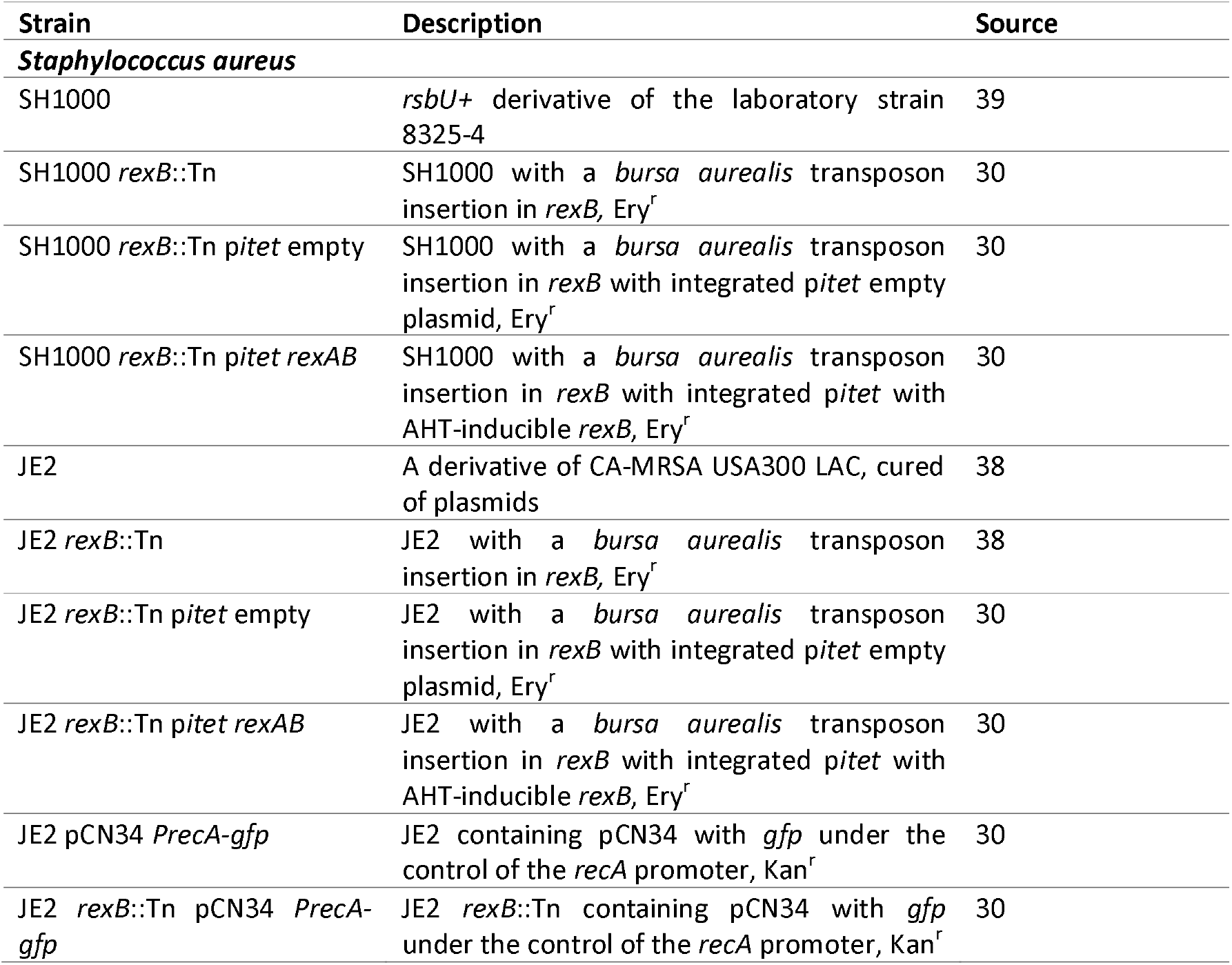
Bacterial strains used in this study.

### *recA-gfp* fluorescent reporter assay

As described previously [30,34], promoter-reporter gene constructs in JE2 and SH1000 backgrounds were used to quantify expression of recA. Antibiotic two-fold dilutions were made in flat-bottomed black-walled 96-well plates containing TSB and inoculated with 1/10 dilution of a stationary phase culture of the reporter strains. Plates were placed into an Infinite M200-PRO microplate reader (Tecan) where cultures were grown for 17 h at 37 °C (700 rpm), and both absorbance at 600 nm (OD_600_) and GFP relative fluorescence units (RFU) were measured every 30 min. To account for differences in cell-density, RFU values were normalised by OD_600_ data at each time point.

### Determination of minimum inhibitory concentration (MIC)

Minimum inhibitory concentrations (MICs) were determined using a serial broth dilution protocol as described previously [30, 58]. Bacteria were diluted to 1 × 10^5^ CFU ml^−1^ and incubated in flat-bottomed 96-well plates with a range of antibiotic concentrations for 17 h at 37 °C under static conditions (aerobic, anaerobic or 5% CO_2_). Media containing daptomycin was supplemented with 1.25 mM CaCl_2_. The MIC was defined as the lowest concentration at which no growth was observed.

### Antibiotic survival assay

Bacteria were adjusted to 10^8^ CFU ml^−1^ in TSB (*S. aureus*) supplemented with antibiotics at 10× MIC. For aerobic incubation, 3 ml of media were inoculated in 30 ml universal tubes and incubated with shaking at 180 rpm. For anaerobic conditions 6 ml of pre-reduced media in 7 ml bijou tubes was inoculated and incubated statically in an anaerobic cabinet. Cultures were incubated at 37 °C and bacterial viability determined by CFU counts. Culture media containing daptomycin was supplemented with 1.25 mM CaCl_2_. Survival was calculated as a percentage of the number of bacteria in the starting inoculum.

### Statistical analyses

Data are represented as the mean or median from three or more independent experiments and analysed by one-way ANOVA corrected for multiple comparison, as described in the figure legends. For each experiment, “n” refers to the number of independent biological replicates. P < 0.05 was considered significant between data points (GraphPad Prism 7 for Windows).

## Acknowledgements

A.M.E. and R.S.C. acknowledge funding from Shionogi & Co., Ltd. A.M.E. also acknowledges support from the National Institute for Health Research (NIHR) Imperial Biomedical Research Centre (BRC). Both authors acknowledge the provision of strains by the Network on Antimicrobial Resistance in Staphylococcus aureus (NARSA) Program: under NIAID/ NIH Contract No. HHSN272200700055C. The funders had no role in the study design, interpretation of the findings or the writing of the manuscript.

