## Supplementary data file for "Multiple classes of bactericidal antibiotics cause DNA double strand breaks in *Staphylococcus aureus*"

Supplementary figures S1

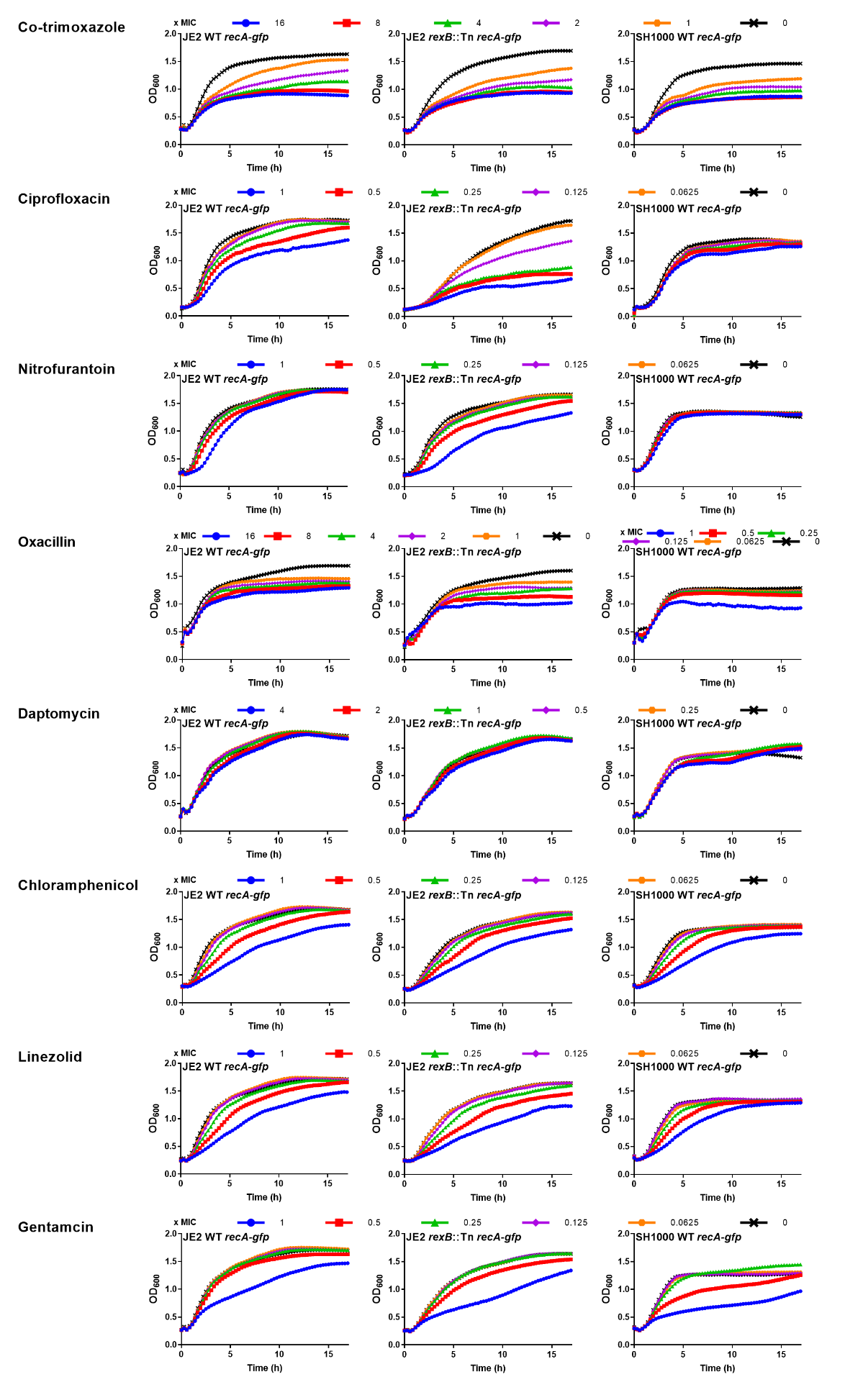

**Figure S1. Representative graphs showing magnitude of growth inhibition of *S. aureus* at concentrations of antibiotics used for Fig. 1 and Fig. 2.** Concentrations of antibiotics are labelled by multiple of the MIC of the individual WT strain (x MIC). Error bars were omitted for clarity.
